# Accounting for genotype uncertainty in the estimation of allele frequencies in autopolyploids

**DOI:** 10.1101/021907

**Authors:** Paul D. Blischak, Laura S. Kubatko, Andrea D. Wolfe

**Author notes:** **Corresponding author:** Paul Blischak, Ohio State University, Dept. of Evolution, Ecology and Organismal Biology, 318 W. 12th Avenue, Columbus, OH 43210.

## Abstract

Despite the increasing opportunity to collect large-scale data sets for population genomic analyses, the use of high throughput sequencing to study populations of polyploids has seen little application. This is due in large part to problems associated with determining allele copy number in the genotypes of polyploid individuals (allelic dosage uncertainty–ADU), which complicates the calculation of important quantities such as allele frequencies. Here we describe a statistical model to estimate biallelic SNP frequencies in a population of autopolyploids using high throughput sequencing data in the form of read counts. We bridge the gap from data collection (using restriction enzyme based techniques [e.g., GBS, RADseq]) to allele frequency estimation in a unified inferential framework using a hierarchical Bayesian model to sum over genotype uncertainty. Simulated data sets were generated under various conditions for tetraploid, hexaploid and octoploid populations to evaluate the model’s performance and to help guide the collection of empirical data. We also provide an implementation of our model in the R package polyfreqs and demonstrate its use with two example analyses that investigate (i) levels of expected and observed heterozygosity and (ii) model adequacy. Our simulations show that the number of individuals sampled from a population has a greater impact on estimation error than sequencing coverage. The example analyses also show that our model and software can be used to make inferences beyond the estimation of allele frequencies for autopolyploids by providing assessments of model adequacy and estimates of heterozygosity.

## Introduction

Biologists have long been fascinated by the occurrence of whole genome duplication (WGD) in natural populations and have recognized its role in the generation of biodiversity (Clausen *et al.* 1940; Stebbins 1950; Grant 1971; Otto & Whitton 2000). Though WGD is thought to have occurred at some point in nearly every major group of eukaryotes, it is a particularly common phenomenon in plants and is regarded by many to be an important factor in plant diversification (Wood *et al.* 2009; Soltis *et al.* 2009; Scarpino *et al.* 2014). The role of polyploidy in plant evolution was originally considered by some to be a “dead-end” (Stebbins 1950; Wagner 1970; Soltis *et al.* 2014) but, since its first discovery in the early twentieth century, polyploidy has been continually studied in nearly all areas of botany (Winge 1917; Winkler 1916; Clausen *et al.* 1945; Grant 1971; Stebbins 1950; Soltis *et al.* 2003, 2010; Soltis & Soltis 2009; Ramsey & Ramsey 2014). Though fewer examples of WGD are currently known for animal systems, groups such as amphibians, fish, and reptiles all exhibit polyploidy (Allendorf & Thorgaard 1984; Gregory & Mable 2005). Ancient genome duplications are also thought to have played an important role in the evolution of both plants and animals, occurring in the lineages preceeding the seed plants, angiosperms and vertebrates (Ohno 1970; Otto & Whitton 2000; Furlong & Holland 2001; Jiao *et al.* 2011). These ancient WGD events during the early history of seed plants and angiosperms have been followed by several more WGDs in all major plant groups (Cui *et al.* 2006; Scarpino *et al.* 2014; Cannon *et al.* 2014). Recent experimental evidence has also demonstrated increased survivorship and adaptability to foreign environments of polyploid taxa when compared with their lower ploidy relatives (Ramsey 2011; Selmecki *et al.* 2015).

Polyploids are generally divided into two types based on how they are formed: auto- and allopolyploids. Autopolyploids form when a WGD event occurs within a single evolutionary lineage and typically have polysomic inheritance. Allopolyploids are formed by hybridization between two separately evolving lineages followed by WGD and are thought to have mostly disomic inheritance. Multivalent chromosome pairing during meiosis can occur in allopolyploids, however, resulting in mixed inheritance patterns across loci in the genome [segmental allopolyploids] (Stebbins 1950). Autopolyploids can also undergo double reduction, a product of multivalent chromosome pairing wherein segments from sister chromatids move together during meiosis—resulting in allelic inheritance that breaks away from a strict pattern of polysomy (Haldane 1930). Autopolyploidy was also thought to be far less common than allopolyploidy, but recent studies have concluded that autopolyploidy occurs much more frequently than originally proposed (Soltis *et al.* 2007; Parisod *et al.* 2010).

The theoretical treatment of population genetic models in polyploids has it origins in the Modern Synthesis with Fisher, Haldane and Wright each contributing to the development of some of the earliest mathematical models for understanding the genetic patterns of inheritance in polyploids (Haldane 1930; Wright 1938; Fisher 1943). Early empirical work on polyploids that influenced Fisher, Haldane and Wright include studies on *Lythrum salicaria* by N. Barlow (1913, 1923), *Dahlia* by W. J. C. Lawrence (1929) and *Primula* by H. J. Muller (1914). The foundation laid down by these early papers has led to the continuing development of population genetic models for polyploids, including models for understanding the rate of loss of genetic diversity and extensions of the coalescent in autotetraploids, as well as modifications of the multispecies coalescent for the inference of species networks containing allotetraploids (Moody *et al.* 1993; Arnold *et al.* 2012; Jones *et al.* 2013). Much of this progress was described in a review by Dufresne *et al.* (2014), who outlined the current state of population genetics in polyploids regarding both molecular techniques and statistical models. Not surprisingly, one of the most promising developments for the future of population genetics in polyploids is the advancement of sequencing technologies. A particularly common method of gathering large data sets for genome scale inferences are restriction enzyme based techniques (e.g., RADseq, ddRAD, GBS, etc.), which we will refer to generally as RADseq (Miller *et al.* 2007; Baird *et al.* 2008; Peterson *et al.* 2012; Puritz *et al.* 2014). However, despite its popularity for population genetic inferences at the diploid level, there are many fewer examples of RADseq experiments conducted on polyploid taxa (but see Ogden *et al.* 2013; Wang *et al.* 2013; Logan-Young *et al.* 2015).

Among the primary reasons for the dearth in applying RADseq to polyploids is the issue of allelic dosage uncertainty (ADU), or the inability to fully determine the genotype of a polyploid organism when it is partially heterozygous at a given locus. This is the same problem that has been encountered by other codominant markers such as microsatellites, which have been commonly used for population genetic analyses in polyploids. One way of dealing with allelic dosage that has been used for multi-allelic microsatellite markers has been to code alleles as either present or absent based on electropherogram readings (allelic phenotypes) and to analyze the resulting dominant data using a program such as polysat (Clark & Jasieniuk 2011; Dufresne *et al.* 2014). de Silva *et al.* (2005) developed a method for inferring allele frequencies using observed allelic phenotype data and used an expectation-maximization algorithm to deal with the incomplete genotype data resulting from ADU. Attempts to directly infer the genotype of polyploid microsatellite loci have also been successfully completed in some cases by using the relative electropherogram peak heights of the alleles in the genotypes (Esselink *et al.* 2004). The estimation problem would be similar for biallelic SNP data collected using RADseq, where a partially heterozygous polyploid will have high throughput sequencing reads containing both alleles. For a tetraploid, the possible genotypes for a partial heterozygote (alleles A and B) would be AAAB, AABB and ABBB. For a hexaploid they are AAAAAB, AAAABB, AAABBB, AABBBB and ABBBBB. In general, the number of possible genotypes for a biallelic locus of a partially heterozygous *K*-ploid (*K* = 3, 4, 5,…) is *K* − 1. A possible solution to this problem for SNPs would be to try to use existing genotype callers and to rely on the relative number of sequencing reads containing the two alleles (similar to what was done for microsatellites). However, this could lead to erroneous inferences when genotypes are simply fixed at point estimates based on read proportions without considering estimation error. Furthermore, when sequencing coverage is low, the number of genotypes that will appear to be equally probable increases with ploidy, making it difficult to distinguish among the possible partially heterozygous genotypes.

In this paper we describe a model that aims to address the problems associated with ADU by treating genotypes as a latent variable in a hierarchical Bayesian model and using high throughput sequencing read counts as data. In this way we preserve the uncertainty that is inherent in polyploid genotypes by inferring a probability distribution across all possible values of the genotype, rather than treating them as being directly observed. This approach has been used by Buerkle & Gompert (2013) to deal with uncertainty in calling genotypes in diploids and the work we present here builds off of their earlier models. Our model assumes that the ploidy level of the population is known and that the genotypes of individuals in the population are drawn from a single underlying allele frequency for each locus. These assumptions imply that alleles in the population are undergoing polysomic inheritance without double reduction, which most closely adheres to the inheritance patterns of an autopolyploid. We acknowledge that the model in its current form is an oversimplification of biological reality and realize that it does not apply to a large portion of polyploid taxa. Nevertheless, we believe that accounting for ADU by modeling genotype uncertainty has the potential to be applied more broadly via modifications of the probability model used for the inheritance of alleles, which could lead to more generalized population genetic models for polyploids (see the **Extensibility** section of the **Discussion**).

## Materials and Methods

Our goal is to estimate the frequency of a reference allele for each locus sampled from a population of known ploidy (*ψ*), where the reference allele can be chosen arbitrarily between the two alleles at a given biallelic SNP. To do this we extend the population genomic models of Buerkle & Gompert (2013), which employ a Bayesian framework to model high throughput sequencing reads (***T***, ***R***), genotypes (***G***) and allele frequencies (***p***), to the case of arbitrary ploidy. The idea behind the model is to view the sequencing reads gathered for an individual as a random sample from the unobserved genotype at each locus. Genotypes can then be treated as a parameter in a probability model that governs how likely it is that we see a particular number of sequencing reads carrying the reference allele. Similarly, we can treat genotypes as a random sample from the underlying allele frequency in the population (assuming Hardy-Weinberg equilibrium). For our model, a genotype is simply a count of the number of reference alleles at a locus which can range from 0 (a homozygote with no reference alleles in the genotype) to *ψ* (a homozygote with only reference alleles in the genotype). All whole numbers in between 0 and *ψ* represent partially heterozygous genotypes. This hierarchical setup addresses the problems associated with ADU by treating genotypes as a latent variable that can be integrated out using Markov chain Monte Carlo (MCMC).

### Model setup

Here we consider a sample of *N* individuals from a single population of ploidy level *ψ* sequenced at *L* unlinked SNPs. The data for the model consist of two matrices containing counts of high throughput sequencing reads mapping to each locus for each individual: ***R*** and ***T***. The *N* × *L* matrix ***T*** contains the total number of reads sampled at each locus for each individual. Similarly, ***R*** is an *N × L* matrix containing the number of sampled reads with the reference allele at each locus for each individual. Then for individual *i* at locus *ℓ,* we model the number of sequencing reads containing the reference allele (*r_iℓ_*) as a Binomial random variable conditional on the total number of sequencing reads (*t_iℓ_*), the underlying genotype (*g_iℓ_*) and a constant level of sequencing error (ϵ)

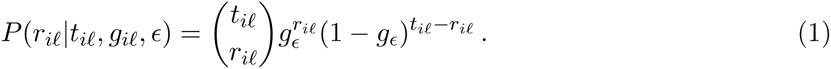

Here *g_ϵ_* is the probability of observing a read containing the reference allele corrected for sequencing error

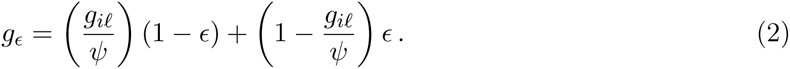

The intuition behind including error is that we want to calculate the probability that we observe a read containing the reference allele. There are two ways that this can happen. (1) Reads are drawn from the reference allele(s) in the genotype with probability 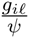 but are only observed as reference reads if they are not errors (probability 1 − *ϵ*). (2) Similarly, reads from the non-reference allele(s) in the genotype are drawn with probability 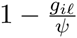 but can be mistakenly read as a coming from a reference allele if an error occurs (probability *ϵ*). The sum across these two possibilities gives the overall probability of observing a read containing the reference allele. If we also assume conditional independence of the sequencing reads given the genotypes, the joint probability distribution for sequencing reads is given by

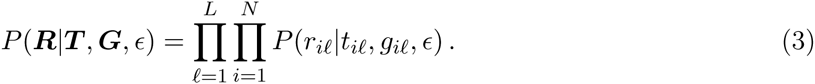

Since the *r_iℓ_*’s are the data that we observe, the product of *P*(*r_iℓ_*|*t_iℓ_*, *g_iℓ_*, *ϵ*) across loci and individuals will form the likelihood in the model.

The next level in the hierarchy is the conditional prior for genotypes. We model each *g_iℓ_* as a Binomial random variable conditional on the ploidy level of the population and the frequency of the reference allele for locus *ℓ* (*p_ℓ_*):

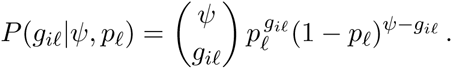

We also assume that the genotypes of the sampled individuals are conditionally independent given the allele frequencies, which is equivalent to taking a random sample from a population in Hardy-Weinberg equilibrium. Factoring the distribution for genotypes and taking the product across loci and individuals gives us the joint probability distribution of genotypes given the ploidy level of the population and the vector of allele frequencies at each locus (*p* = {*p*_1_,…,*p_L_*}):

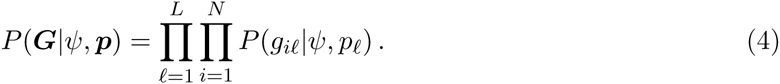

We choose here to ignore other factors that may be influencing the distribution of genotypes such as double reduction. In general, double reduction will act to increase homozygosity (Hardy 2015). However, it is more prevalent for loci that are farther away from the centromere, which makes the estimation of a global double reduction parameter (typically denoted α) inappropriate for the thousands of loci gathered from across the genome using techniques such as RADseq. It might be possible to estimate a per locus rate of double reduction (*α_ℓ_*) but this would add an additional parameter that would need to be estimated for each locus, perhaps unnecessarily if the majority end up being equal, or close, to 0.

The final level of the model is the prior distribution on allele frequencies. Assuming *a priori* independence across loci, we use a Beta distribution with parameters *α* and *β* both equal to 1 as our prior distribution for each locus. A Beta(1,1) is equivalent to a Uniform distribution over the interval [0,1], making our choice of prior uninformative. The joint posterior distribution of allele frequencies and genotypes is then equal to the product across all loci and all individuals of the likelihood, the conditional prior on genotypes and the prior distribution on allele frequencies up to a constant of proportionality

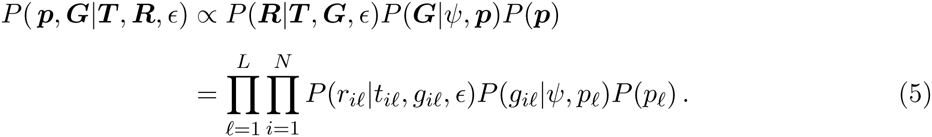

The marginal posterior distribution for allele frequencies can be obtained by summing over genotypes

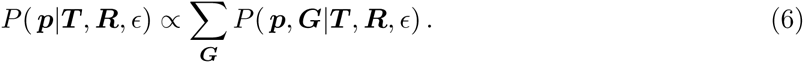

It would also be possible to examine the marginal posterior distribution of genotypes but here we will focus primarily on allele frequencies.

### Full conditionals and MCMC using Gibbs sampling

We estimate the joint posterior distribution for allele frequencies and genotypes in Eq. 5 using MCMC. This is done using Gibbs sampling of the states (***p***, ***G***) in a Markov chain by alternating samples from the full conditional distributions of ***p*** and ***G***. Given the setup for our model using Binomial and Beta distributions (which form a conjugate family), analytical solutions for these distributions can be readily acquired (Gelman *et al.* 2014). The full conditional distribution for allele frequencies is Beta distributed and is given by Eq. 7 below:

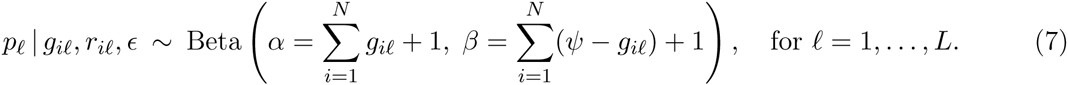

This full conditional distribution for *p_ℓ_* has a natural interpretation as it is roughly centered at the proportion of sampled alleles carrying the reference allele divided by the total number of alleles sampled. The “+1” comes from the prior distribution and will not have a strong influence on the posterior when the sample size is large.

The full conditional distribution for genotypes is a discrete categorical distribution over the possible values for the genotypes (0,…,*ψ*). The distribution for individual *i* at locus *ℓ* is

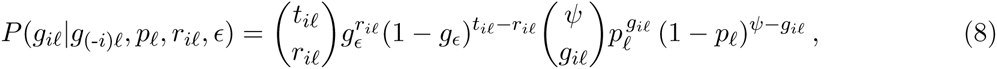

where *g*(*-i*)*ℓ* is the value of the genotypes for all sampled individuals excluding individual *i* and *g_ϵ_* is the same as Eq. 2. The full conditional distribution for genotypes can be seen as the product of two quantities: (1) the probability of each of the possible genotypes based on the observed reference reads and (2) the probability of drawing each genotype given the allele frequency for that locus in the population.

We begin our Gibbs sampling algorithm in a random position in parameter space through the use of uniform probability distributions. The genotype matrix is initialized with random draws from a Discrete Uniform distribution ranging from 0 to *ψ* and the initial allele frequencies are drawn from a Uniform distribution on the interval [0,1].

### Simulation study

Simulations were performed to assess error rates in allele frequency estimation for tetraploid, hexaploid and octoploid populations (*ψ* = 4, 6 and 8, respectively). Data were generated under the model by sampling genotypes from a Binomial distribution conditional on a fixed, known allele frequency (*p_ℓ_* = 0.01, 0.05, 0.1, 0.2, 0.4). Total read counts were simulated for a single locus using a Poisson distribution with mean coverage equal to 5, 10, 20, 50 or 100 reads per individual. We then sampled the number of sequencing reads containing the reference allele from a Binomial distribution conditional on the number of total reads, the genotype and sequencing error (Eq. 1; *ϵ* fixed to 0.01). Finally, we varied the number of individuals sampled per population (*N* = 5, 10, 20, 30) and ran all possible combinations of the simulation settings. Our choice for the number of individuals to simulate was intended to reflect sampling within a *single* population/locality and not that of an entire population genetics study. Furthermore, RAD sequencing is used at various taxonomic levels from population genetics to phylogenetics (e.g., Rheindt *et al.* 2014; Eaton *et al.* 2015), and we wanted our simulations to be informative across these applications. Each combination of sequencing coverage, individuals sampled and allele frequency was analyzed using 100 replicates for tetraploid, hexaploid and octoploid populations for a total of 30,000 simulation runs. MCMC analyses using Gibbs sampling were run for 100,000 generations with parameter values stored every 100th generation. The first 25% of the sample was discarded as burn-in, resulting in 750 posterior samples for each replicate. Convergence on the stationary distribution, *P*(***p***, ***G****|****R****, ϵ*), was assessed by examining trace plots for a subset of runs for each combination of settings and ensuring that the effective sample sizes (ESS) were greater than 200. Deviations from the known underlying allele frequency used to simulate each data set were assessed by taking the posterior mean of each replicate and calculating the root mean squared error (RMSE) based on the true underlying value. We also compared the posterior mean as an estimate of the allele frequency at a locus to a more simple estimate calculated directly from the read counts (mean read ratio): 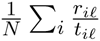. Comparisons between estimates were again made using the RMSE.

All simulations were performed using the R statistical programming language (R Core Team 2014) on the Oakley cluster at the Ohio Supercomputer Center (https://osc.edu). Figures were generated using the R packages ggplot2 (Wickham 2009) and reshape (Wickham 2007), with additional figure manipulation completed using Inkscape (https://inkscape.org). MCMC diagnostics were done using the coda package (Plummer *et al.* 2006). All scripts are available on GitHub (https://github.com/pblischak/polyfreqs-ms-data) in the `‘code/’` folder and all simulated data sets are in the `‘raw_data/’` folder.

### Example analyses of autotetraploid potato (*Solanum tuberosum*)

To further evaluate the model and to demonstrate its use we present an example analysis using an empirical data set collected for autotetraploid potato (*Solanum tuberosum*) using the Illumina GoldenGate platform (Anithakumari *et al.* 2010; Voorrips *et al.* 2011). Though these data aren’t the typical reads returned by RADseq experiments, they still represent the same type of binary response data that our model uses to get a probability distribution for biallelic SNP genotypes. A detailed walkthrough with the code used for each step is provided as Supplemental Material. The data set and output are also available on GitHub (https://github.com/pblischak/polyfreqs-ms-data) in the `‘example/’` folder.

#### Calculating expected and observed heterozygosity

One advantage of using a Bayesian framework for our model is that we can approximate a posterior distribution for any quantity that is a functional transformation of the parameters that we are estimating without doing any additional MCMC simulation (Gelman *et al.* 2014). Two such quantities that are often used in population genetics are the observed and expected heterozygosity, which are in turn used for calculating the various fixation indices (*F_IS_, F_IT_, F_ST_*) introduced by Wright (1951). To analyze levels of heterozygosity in this way, we used the estimators of Hardy (2015) to calculate the per locus observed (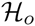) and expected (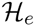) heterozygosity for each stored sample of the joint posterior distribution in Eq. 5. This procedure is especially useful because it estimates heterozygosity while taking into account ADU by utilizing the marginal posterior distribution of genotypes. Given a total of *M* posterior samples of genotypes and allele frequencies, we calculate the *m*^th^ (*m* = 1,…, *M*) estimate of the observed heterozygosity using Eq. 9 [numerator of Eq. 7 in Hardy (2015)]:

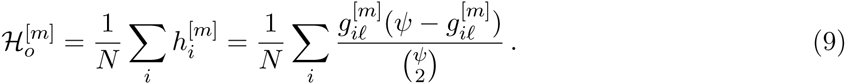

Similarly, the *m*^th^ estimate of the expected heterozygosity is calculated using Eq. 10 [denominator of Eq. 8 in Hardy (2015)]:

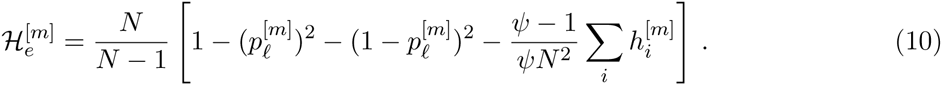

The posterior distribution of a multi-locus estimate of heterozygosity can then be approximated by taking the average across loci for each of the per locus posterior samples.

To evaluate levels of heterozygosity in autotetraploid potato, we obtained biallelic count data for 224 accessions collected at 384 loci using the Illumina GoldenGate platform from the R package fitTetra (Voorrips *et al.* 2011), which provides the data set as part of the package. We chose the ‘X’ reading to be the count data for the reference allele and added the ‘X’ and ‘Y’ readings together to get the total read counts (‘X’ and ‘Y’ represent the counts of the two alternative alleles). Initial attempts to analyze the data set using our Gibbs sampling algorithm were unsuccessful due to arithmetic underflow. This was due to the fact that the counts/intensities returned by the Illumina GoldenGate platform are on a different scale (~10,000−20,000+) than the read counts that would be expected from a RADseq experiment. To alleviate this problem, we rescaled the data set while preserving the relative dosage information by dividing the GoldenGate count readings by 100 and rounding to the nearest whole number. We then analyzed the rescaled count data using 100,000 MCMC generations, sampling every 100 generations and using the stored samples of the allele frequencies and genotypes to calculate the observed and expected heterozygosity for a total of 1,000 posterior samples of the per locus observed and expected heterozygosity. We also compared post burn-in (25%) allele frequency estimates based on the posterior mean to the simple allele frequency estimate based directly on read counts used previously (mean read ratio). Posterior distributions for multi-locus estimates of observed and expected heterozygosity were obtained by taking the average across loci for each posterior sample of the per locus estimates using a burn-in of 25%.

#### Evaluating model adequacy

As noted earlier, the probability model that we use for the inheritance of alleles is one of polysomy without double reduction. In some cases, this model may be inappropriate. Therefore, it can be informative to check for loci that do not follow the model that we assume. Below we describe a procedure for rejecting our model of inheritance on a per locus basis using comparisons with the posterior predictive distribution of sequencing reads. Model checking is an important part of making statistical inferences and can play a role in understanding when a model adequately describes the data being analyzed. In the case of our model, it can serve as a basis for understanding the inheritance patterns of the organism being studied by determining which loci adhere to a simple pattern of polysomic inheritance. Other sources of disequilibrium that could indicate poor model fit include inbreeding, null alleles and allele drop out (*sensu* Arnold *et al.* 2013), making this posterior predictive model check more broadly applicable for RADseq data.

Given *M* posterior samples for the allele frequencies at locus 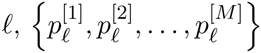 we simulate new values for the genotypes 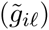 and reference read counts 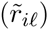 for all individuals and use the ratio of simulated reference read counts to observed total read counts 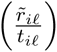 as a summary statistic for comparing the observed read count ratios to the distribution of the predicted read count ratios. The use of the likelihood (or similar quantities) as a summary statistic has been a common practice in posterior predictive comparisons of nucleotide substitution models, and more recently for comparative phylogenetics (Ripplinger & Sullivan 2010; Reid *et al.* 2014; Pennell *et al.* 2015). We use the ratio of reference to total read counts here because it is the maximum likelihood estimate of the probability of success for a Binomial random variable and because it is a simple quantity to calculate. The use of other summary statistics, or a combination of multiple summary statistics, would also possible. The procedure for our posterior predictive model check is as follows:

1. For locus *ℓ* = 1,…, *L*:

1.1. For posterior sample *m* = 1,…, *M*:

1.1.1. Simulate new genotype values 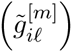 for all individuals (*i* = 1,…, *N*) by drawing from a Binomial 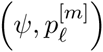.
1.1.2. Simulate new reference read counts 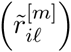 from each new genotype for all individuals by drawing from Eq. 1.
1.1.3. Calculate the reference read ratio for the simulated data for sample *m* and sum across individuals: 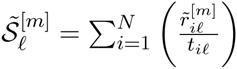
1.1.4. Calculate the reference read ratio for the observed data and sum across individuals: 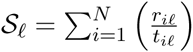
1.2. Calculate the difference between the observed reference read ratio and the *M* simulated reference read ratios: 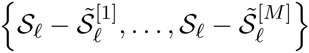
2. Determine if the 95% highest posterior density (HPD) interval of the distribution of re-centered reference read ratios contains 0.

When the distribution of the differences in ratios between the observed and simulated data sets does not contain 0 in the 95% HPD interval, it provides evidence that the locus being examined does not follow a pattern of strict polysomic inheritance. A similar approach could be used on an individual basis by comparing the observed ratio of reference reads to the predicted ratios for each individual at each locus. We used this posterior predictive model checking procedure to assess model adequacy in the potato data set using the posterior distribution of allele frequencies estimated in the previous section with 25% of the samples discarded as burn-in.

## Results

Our Gibbs sampling algorithm was able to accurately estimate allele frequencies for a number of simulation settings while simultaneously allowing for genotype uncertainty. There were no indications of a lack of convergence (ESS values > 200) for any of the simulation replicates and all trace plots examined also indicated that the Markov chain had reached stationarity. Running the MCMC for 100,000 generations and sampling every 100th generation appeared to be suitable for our analyses and we recommend it as a starting point for running most data sets. Reducing the number of generations and sampling more frequently (e.g., 50,000 generations sampled every 50 generations) could be a potential work around for larger data sets. When doing test runs we went as low as 20,000 generations sampled every 20th generation, which still passed our diagnostic tests for convergence. This is likely because the parameter space of our model is not overly difficult to navigate so stationarity is reached rather quickly. Ultimately, the deciding factor on how long to run the analysis and how frequently to sample the chain will come down to assessing convergence.

### Simulation study

Increasing the number of individuals sampled had the largest effect on the accuracy of allele frequency estimation (Figure 1). Since allele frequencies are population parameters, it is not surprising that sampling more individuals from the population leads to better estimates. This appears to be the case even when sequencing coverage is quite low (5x, 10x), which corroborates the observations made by Buerkle & Gompert (2013). This is not to say, however, that sequencing coverage has no effect on the posterior distribution of allele frequencies. Lower sequencing coverage affects the posterior distribution by increasing the posterior standard deviation (Figure 2). An interesting pattern that emerged during the simulation study is the observation that the allele frequencies closer to 0.5 tend to have higher error rates, which is to be expected given that the variance of a Binomial random variable is highest when the probability of success is 0.5. We also observed small differences in the RMSE between ploidy levels, with estimates increasing in accuracy with increasing ploidy. Comparisons between the posterior mean and mean read ratio estimates of allele frequencies (Figure S1) show that the estimate based on read ratios has a lower RMSE than the posterior mean when the true allele frequency is low (*p_ℓ_* = 0.01, 0.05) but has higher error rates than the posterior mean for allele frequencies closer to 0.5. When sequencing coverage is greater than 10x and the number of individuals sampled is greater than 20, the two estimates are almost indistinguishable.

**Figure 1:**
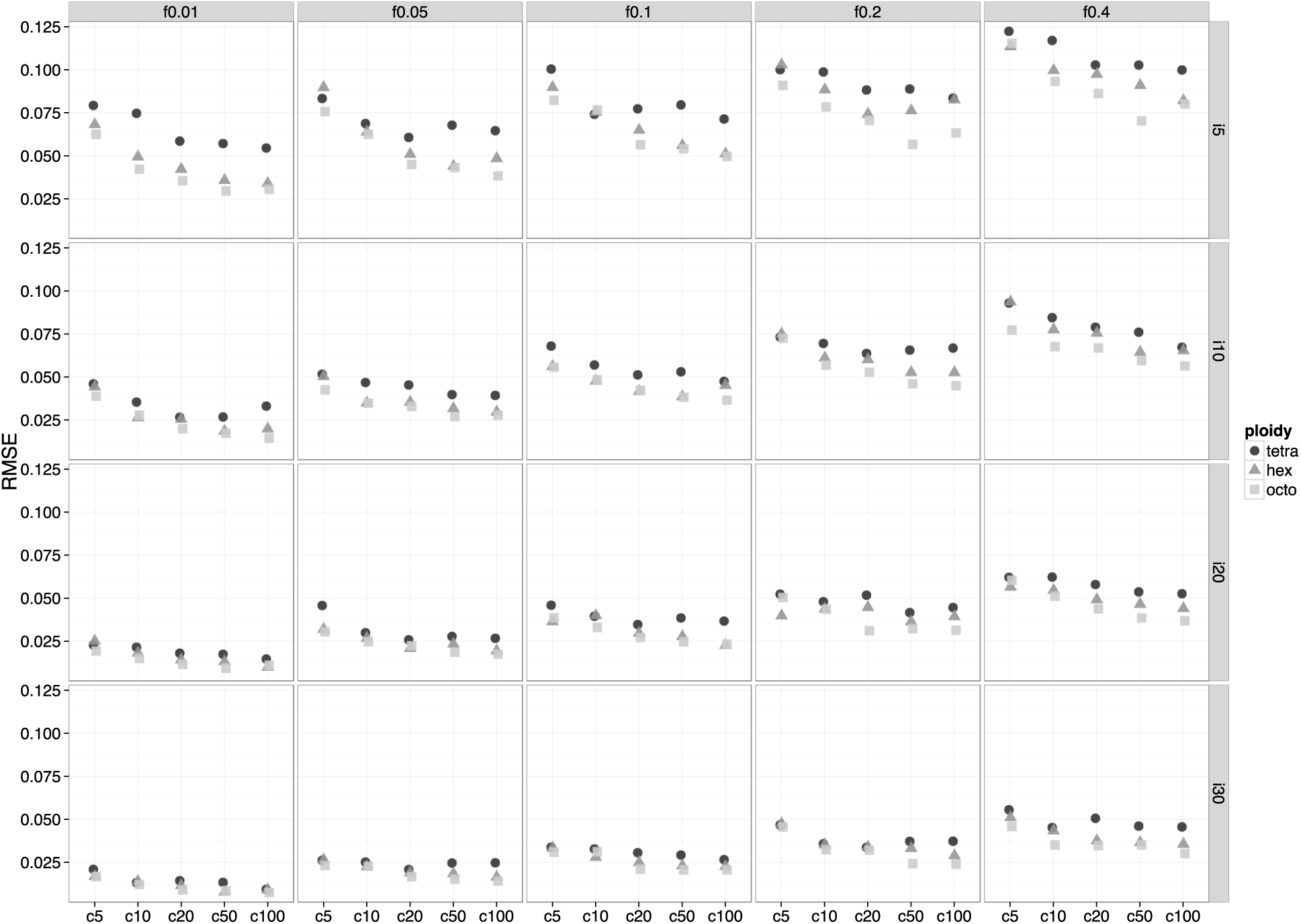
Error in allele frequency estimation as measured by the RMSE of posterior means. Columns represent the different allele frequencies used to simulate read data (0.01, 0.05, 0.1, 0.2, 0.4), rows represent the number of individuals samples from the population (5, 10, 20, 30). Each individual plot shows the RMSE of the estimates for each ploidy level (tetra, hex, octo) across the different levels of coverage (5x, 10x, 20x, 50x 100x). The best scenario is in the bottom left with 30 individuals sampled and an allele frequency of 0.01. The worst scenario is in the upper right corner with 5 individuals sampled and an allele frequency of 0.4. Looking across rows shows that error increases as allele frequencies get closer to 0.5. Looking up and down columns shows that error increases as the number of individuals decreases. Within each plot, increasing sequence coverage does not have as large of an effect on error, and differences in ploidy show that error decreases as ploidy increases.

**Figure 2:**
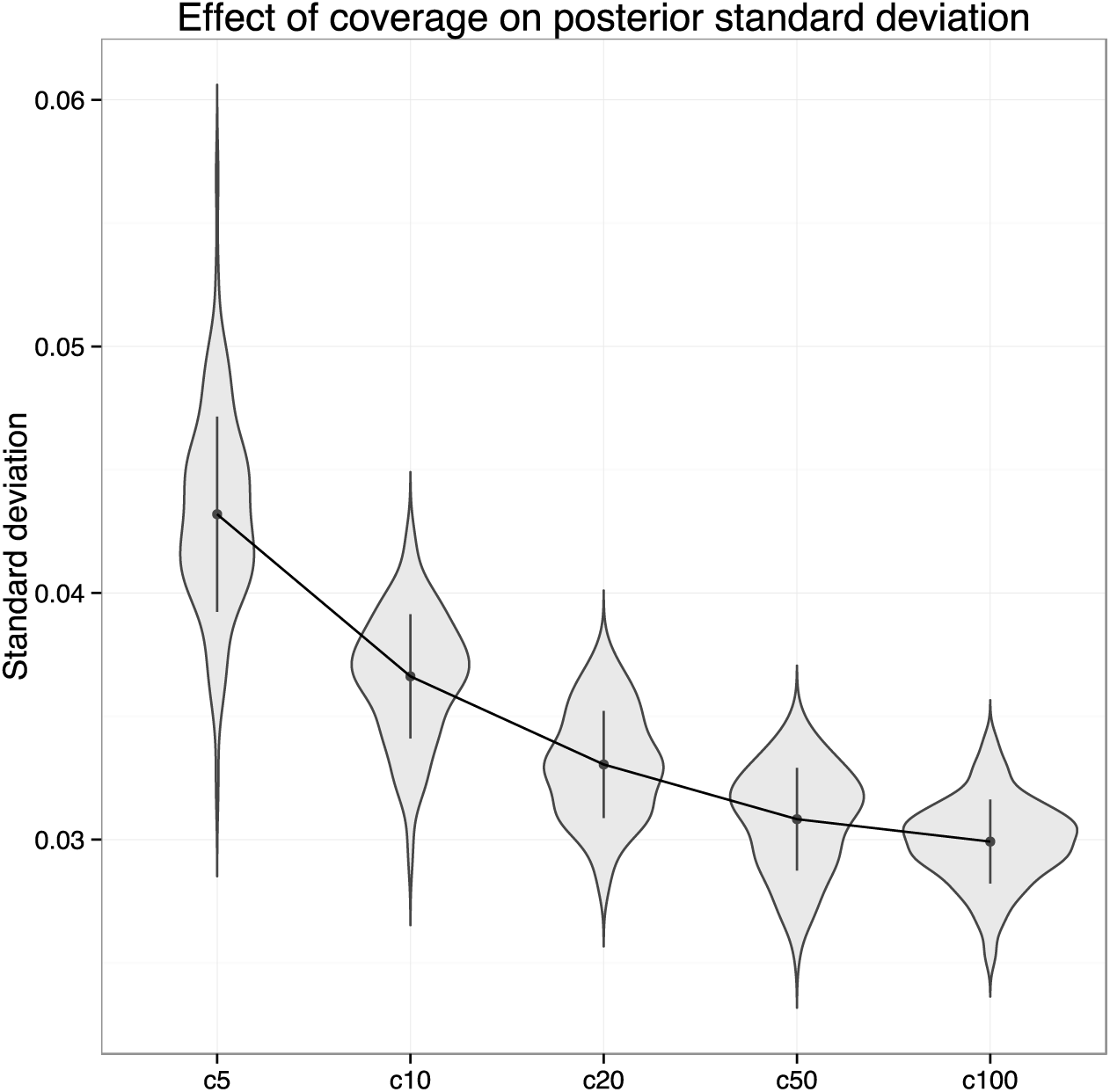
The posterior standard deviation for allele frequencies decreases compared across levels of sequencing coverage. This plot provides a comparison of the distribution of the posterior standard deviations of the 100 replicates performed for each level of sequencing coverage (5x, 10x, 20x, 50x, 100x) for the hexaploid simulation with 30 individuals sampled from the population and an allele frequency of 0.2.

### Example analyses

Our analyses of *Solanum tuberosum* tetraploids showed levels of heterozygosity consistent with a pattern of excess outbreeding 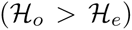. In fact, the posterior distributions of the multi-locus estimates of observed and expected heterozygosity do not overlap at all (Figure 3). The assessment of model adequacy also showed that 49 out of the 384 loci (~13%) were a poor fit to the model of polysomic inheritance that we assume. The allele frequency estimates using the posterior mean and the mean read ratio provided similar estimates and were comparable for most loci. For loci in which the frequency of the reference allele is very low, the read ratio estimate tends to be higher than the posterior mean. However, the overall pattern does not indicate over or under estimation for most allele frequencies (Figure S2). When we took the difference between the estimates at each locus, the distribution was centered near 0 (Figure S3).

**Figure 3:**
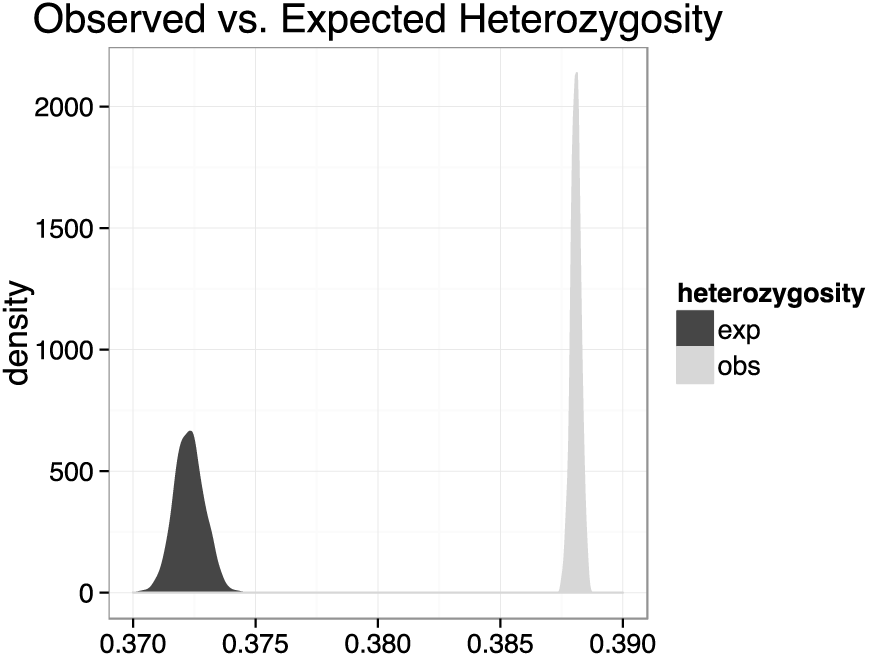
Posterior distributions of the multi-locus estimates of expected and observed heterozygosity in *Solanum tuberosum.* The observed heterozygosity is higher than the expected, consistent with a pattern of excess outbreeding.

## Discussion

The inference of population genetic parameters and the demographic history of non-model polyploid organisms has consistently lagged behind that of diploids. The difficulties associated with these inferences present themselves at two levels. The first of these is the widely known inability to determine the genotypes of polyploids due to ADU. Even though there have been theoretical developments in the description of models for polyploid taxa as early as the 1930s, a large portion of this population genetic theory relies on knowledge about individuals’ genotypes (e.g., Haldane 1930; Wright 1938). The second complicating factor is the complexity of inheritance patterns and changes in mating systems that often accompany WGD events. Polyploid organisms can sometimes mate by both outcrossing or selfing, and can display mixed inheritance patterns at different loci in the genome (Dufresne *et al.* 2014). If genotypes were known, then it might be easier to develop and test models for dealing with and inferring rates of selfing versus outcrossing, as well as understanding inheritance patterns across the genome. However, ADU only compounds the problems associated with these inferences, making the development and application of appropriate models far more difficult (but see list of software in Dufresne *et al.* 2014). The model we have presented here deals with the first of these two issues by not treating genotypes as observed quantities. Almost all other methods of genotype estimation for polyploids treat the genotype as the primary parameter of interest. Our model is different in that we still use the read counts generated by high throughput sequencing platforms as our observed data but instead integrate across genotype uncertainty when inferring other parameters, thus bypassing the problems caused by ADU.

Despite our focus on bypassing ADU, an important consideration for the model we present here is that, because it approximates the joint posterior distribution of allele frequencies and genotypes, it would also be possible to use the marginal posterior distribution of genotypes to make inferences using existing methods. This could be done using the posterior mode as a maximum *a posteriori* (MAP) estimate of the genotype for downstream analyses, followed by analyzing the samples taken from the marginal posterior distribution of genotypes. The resulting set of estimates would not constitute a “true” posterior distribution of downstream parameters but would allow researchers to interpret their results based on the MAP estimate of the genotypes while still getting a sense for the amount of variation in their estimates. Using the marginal posterior distribution of genotypes in this way could technically be applied to any type of polyploid, but is only really appropriate for autopolyploids due to the model of inheritance that is used. Other methods for estimating SNP genotypes from high throughput sequencing data include the program SuperMASSA, which models the relative intensity of the two alternative alleles using Normal densities (Serang *et al.* 2012).

A second important factor for using our model is that, although estimates of allele frequencies can be accurate when sequencing coverage is low and sample sizes are large (see Figure S4 for a direct comparison between sample size and coverage), the resulting distribution for genotypes is likely going to be quite diffuse. For analyses that treat genotypes as a nuisance parameter, this is not an issue since we can integrate across genotype uncertainty. However, if the genotype *is* of primary interest, then the experimental design of the study will need to change to acquire higher coverage at each locus for more accurate genotype estimation. Therefore, the decision between sequencing more individuals with lower average coverage versus sequencing fewer individuals with higher average coverage depends primarily on whether the genotypes will be used or not.

### Extensibility

The modular nature of our hierarchical model can allow for the addition and modification of levels in the hierarchy. One of the simplest extensions to the model that can build directly on the current setup would be to consider loci with more than two alleles. This can be done using Multinomial distributions for sequencing reads and genotypes and a Dirichlet prior on allele frequencies (the Multinomial and Dirichlet distributions form a conjugate family; Gelman *et al.* 2014). We could also model populations of mixed ploidy by using a vector of individually assigned ploidy levels instead of assuming a single value for the whole population (*ψ* = {*ψ*_1_,…,*ψ_N_*}). However, this would assume random mating among ploidy levels.

#### Double reduction

The inclusion of double reduction into the model is a difficult consideration for genome wide data collected using high throughput sequencing platforms. The number of parameters estimated by our model is *L* × (*N* + 1) and including double reduction would add an additional *L* parameters, bringing the total to *L* × (*N* + 2). Though the addition of these parameters would not prohibit an analysis using Gibbs sampling, we chose to implement the simpler equilibrium model. We hope to include double reduction in future models but feel that our posterior predictive model checking procedure will prove sufficient for identifying loci in disequilibrium with our current implementation. Another concern that we had regarding double reduction is that it can be confounded with the overall signal of inbreeding, making it especially difficult to tease apart the specific effects of double reduction alone (Hardy 2015). However, because the probability of double reduction at a locus (*α_ℓ_*) depends on its distance from the centromere (call it *x_ℓ_*), a potential way to estimate *α_ℓ_* would be to use the *α_ℓ_*’s as predictor variables in a linear model: *α_ℓ_* = *β*_0_ + *β*_1_*x_ℓ_*. This would only add two additional parameters (*β*_0_ and *β*_1_) that would need to be estimated and would be completely independent of the number of loci analyzed. The downside to this approach is that it would only be applicable for polyploid organisms with sequenced genomes (or the genome of a diploid progenitor), making the use of such a model impractical for the time being.

#### Additional levels in the hierarchical model

The place where we believe our model could have the greatest impact is through modifications and extensions of the probability model used for the inheritance of alleles. These models have been difficult to apply in the past as a result of genotype uncertainty. However, using our model as a starting point, it could be possible to infer patterns of inheritance (polysomy, disomy, heterosomy) and other demographic parameters (e.g., effective population size, population differentiation) without requiring direct knowledge about the genotypes of the individuals in the population. For example, Haldane’s (1930) model of genotype frequencies for autopolyploids that are partially selfing could be used to infer the prevalence of self-fertilization within a population. Another possible approach would be to use general disequilibrium coefficients (*D_A_*) to model departures from Hardy-Weinberg equilibrium (Hernández & Weir 1989; Weir 1996). A more recent model described by Stift *et al.* (2008) used microsatellites to infer the different inheritance patterns (disomic, tetrasomic, intermediate) for tetraploids in the genus *Rorippa* (Brassicaceae) following crossing experiments. The reformulation of such a model for biallelic SNPs gathered using high throughput sequencing could provide a suitable framework for understanding inheritance patterns across the genome. An ideal model would be one that could help to understand genome-wide inheritance patterns for a polyploid of arbitrary formation pathway (autopolyploid ↔ allopolyploid) without the need conduct additional experiments. However, to our knowledge, such a model does not currently exist.

## Conclusions

The recent emergence of models for genotype uncertainty in diploids has introduced a theoretical framework for dealing with the fact that genotypes are unobserved quantities (Gompert & Buerkle 2012; Buerkle & Gompert 2013). Our extension of this theory to cases of higher ploidy (specifically to autopolyploids) progresses naturally from the original work but also serves to alleviate the deeper issue of ADU. The power and flexibility of these models as applied at the diploid level has the potential to be replicated for polyploid organisms with the addition of suitable models for allelic inheritance. The construction of hierarchical models containing probability models for ADU, allelic inheritance and perhaps even additional levels for important parameters such as F-statistics or the allele frequency spectrum also have the potential to provide key insights into the population genetics of polyploids (Gompert & Buerkle 2011; Buerkle & Gompert 2013). Future work on such models will help to progress the study of polyploid taxa and could eventually lead to more generalized models for understanding the processes that have shaped their evolutionary histories.

## Software note

We have combined the scripts for our Gibbs sampler as an R package—polyfreqs—which is available on GitHub (https://github.com/pblischak/polyfreqs). Though polyfreqs is written in R, it deals with the large data sets that are generated by high throughput sequencing platforms in two ways. First, it takes advantage of R’s ability to incorporate C++ code via the Rcpp and RoppArmadillo packages, allowing for a faster implementation of our MCMC algorithm (Eddelbuettel & François 2011; Eddelbuettel 2013; Eddelbuettel & Sanderson 2014). Second, since the model assumes independence between loci, polyfreqs can facilitate the process of parallelizing analyses by splitting the total read count and reference read count matrices into subsets of loci which can be analyzed at the same time on separate nodes of a computing cluster. Additional features of the program include:

- Estimation of posterior distributions of per locus observed and expected heterozygosity (`het_obs` and `het_exp`, respectively).
- Maximum *a posteriori* (posterior mode) estimation of genotypes using the `get_map_genotypes()` function.
- Posterior predictive model checking using the `polyfreqs_pps()` function.
- Simulation of high throughput sequencing read counts and genotypes from user specified allele frequencies using the `sim_reads()` function.
- Options for controlling program output such as writing genotype samples to file, printing MCMC updates to the R console, etc.
- Simple input format using tab delimited text files that can be directly imported into R using the `read.table()` function. The format is as follows:

1. An optional row of locus names (use `header=TRUE` to specify this in `read.table()`).
2. One row for each individual.
3. First column contains individual names (use `row.names=1` to specify this in `read.table()`).
4. One column for each locus.

## Acknowledgements

The authors would like to thank the Ohio Supercomputer Center for access to computing resources and Nick Skomrock for assistance with deriving the full conditional distributions of the model in the diploid case. We would also like to thank Frederic Austerlitz, Aaron Wenzel, members of the Wolfe and Kubatko labs and 3 anonymous reviewers for their helpful comments on the manuscript. This work was partially funded through a grant from the National Science Foundation (DEB-1455399) to ADW and LSK.

## Author Contributions

Conceived of the study: PDB, LSK and ADW. PDB derived the polyploid model, ran the simulations and other analyses, coded the R package and wrote the initial draft of the manuscript. PDB, LSK and ADW reviewed all parts of the manuscript and all authors approved of the final version.

## Data Accessibility

Scripts for simulating the data sets, analyzing them using Gibbs sampling and producing the figures from the resulting output can all be found on GitHub, along with the original simulated data sets and autotetraploid potato data (https://github.com/pblischak/polyfreqs-ms-data). We also provide an implementation of the Gibbs sampler for estimating allele frequencies in the R package polyfreqs (https://github.com/pblischak/polyfreqs). See the package vignette or GitHub wiki for more details (https://github.com/pblischak/polyfreqs/wiki).

**Table 1:**
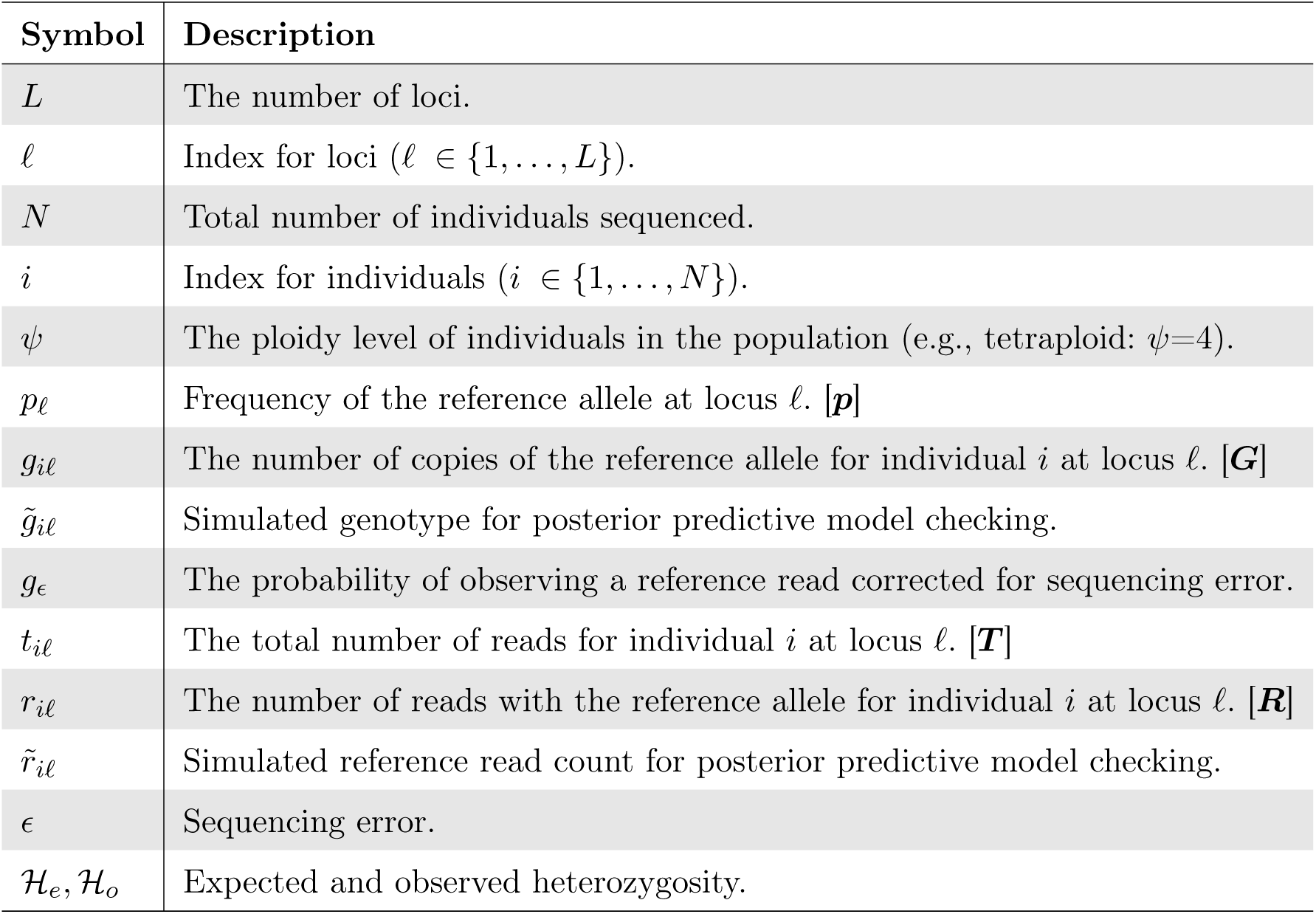
Notation and symbols used in the description of the model for estimating allele frequencies in polyploids. Vector and matrix forms of the variables are also provided when appropriate.

